# Contrasted effect of climate and anthropogenic change on future invasion risk of a solitary bee *Amegilla pulchra*

**DOI:** 10.1101/2024.12.20.629635

**Authors:** Nicolas Dubos, Benoit Geslin, Hervé Jourdan, David Renault, Marie Zakardjian

**Author notes:** Corresponding author: Nicolas Dubos.

## Abstract

*Amegilla pulchra* is a solitary bee from Australia that has recently been spread throughout many islands of the Pacific. The non-regulated human-driven spread of the species may affect the local pollinator communities and their interactions with host plants. We used an ecological niche modelling approach, accounting for non-equilibrium and anthropogenic spread with the most recently recommended approach, and predicted the potential spread of the species under current and future conditions. We expected climate change and increase in human density to offer new suitable environments for the spread of the species. Invasion risks will increase in the future overall, but more in the non-native regions compared to the native region. In the native region, the projected effect of future environmental change was highly contrasted, with increasing invasion risk in human-dense areas but decreasing elsewhere. We found high risks of invasion in eastern Asia and provided a world ranking of entry points for surveillance priority which accounts for maritime traffic. This study highlights potential contrasted effects between climate and anthropogenic change, with differing projections between the native and the non-native regions. Public awareness and prevention will be the key to prevent further spread and mitigate potential adverse effects of the species on island systems. In regions that are already invaded, we propose that habitat restoration is a promising strategy for both the mitigation of the spread and the conservation of local communities.

## 1. Introduction

Invasion risks are not homogeneous through space nor time. The regions at risk depend on how suitable are the local environmental conditions to a given invasive alien species (IAS), and how anthropogenic activity can facilitate the introductions and spread of non-native species (Bellard et al., 2016b). The risk may also vary in either direction with temporal change in the environment— such as seasonal meteorological variations, climate change, or human-induced alterations of habitats (Bellard et al., 2013). The consequences of biological invasions include biodiversity loss and high economic costs related to damage and management (Diagne et al., 2021), and may severely affect pollination service (IPBES, 2016). The risks are particularly high on islands where the rate of endemism is often higher in comparison to continental ecosystems (Bellard et al., 2016a). However, the climate may change differently between continents and islands (IPCC, 2021). Yet, predictions of future invasion risks are usually quantified globally and comparisons between the continent (i.e. often corresponding to the native region; Luque et al., 2014) and invaded islands are rarely documented.

Prevention is the best strategy to mitigate the ecological and economic impacts incurred by IAS. It is crucial to promote proactive surveillance and early detection in the regions the most at risk and at potential entry points (Cuthbert et al., 2022). The regions the most at risk in the present and future can be identified with predictive modelling approaches such as Species Distribution Models (SDMs; also called Environmental Niche Models, ENMs). These models are increasingly used to predict species response to future environmental change (Dubos et al., 2023b), prioritizing conservation areas (Dubos et al., 2022b; Leroy et al., 2014), testing ecological hypotheses (Anderson et al., 2009; Raxworthy et al., 2007) or assessing invasion risk (Gallien et al., 2012). However, the necessary complexity of their design, together with their increasing accessibility with user-friendly interfaces has led to a widespread misuse or misinterpretation of SDMs (Leroy, 2023; Yackulic et al., 2013; Zurell et al., 2020). As a result, projections of future invasion risk can largely differ and become misleading. Provided they are carefully designed, they can produce maps that help targeting priority entry points for stakeholders (Hui, 2022).

Modelling the environmental niche of non-native species that are still spreading is challenging because of the non-equilibrium hypothesis (Gallien et al., 2012; Hui, 2022). After being introduced outside of their native range, IAS usually extent their realized niche as a result of ecological release (i.e. fewer biotic constrains) and in situ adaptation (Fieldsend et al., 2021). At an early stage of range expansion, population sizes are still small and hard to detect, hence the available occurrence data does not represent well the potential niche to be filled in the future. Therefore, models might downplay the importance of some suitable regions that have not been invaded yet and underestimate the potential spread. To mitigate this effect, it is recommended to include both data from the native and non-native range to train the models with all the available information (Hui, 2022). In addition, removing the zones that have not been reached by the species from the model training area (i.e. background) might reduce omission errors and the effect of non-equilibrium. To do so, the model training area should be limited to a restricted area near the known occurrences, and give more weight to the most visited places (i.e. the equivalent of a sample bias correction; Phillips et al., 2009a). Eventually, since the non-native species could be introduced into places that are spatially separated, it is of utmost importance to evaluate model spatial transferability with appropriate approaches (e.g. block-cross validation; Valavi et al., 2023).

To date, 80 species of wild bees have been detected outside of their native range. Despite being overlooked, some of them have already impacted native ecosystems (Geslin et al., 2023). Alien bees can alter native pollinators through several mechanisms, including competition for nesting sites and floral resources, transmission of diseases and pathogens (Morales et al., 2013; Zakardjian et al., 2023a). Moreover, by visiting local plant communities, alien bees may reduce native plants reproductive success and promote alien hosts (Aizen et al., 2014; Chalcoff et al., 2022; Dohzono and Yokoyama, 2010). The spread might represent a threat to native pollinators (Zakardjian et al., 2023a) and plant-pollinator networks (Zakardjian et al., 2023b), potentially leading to co-extinctions and loss of phylogenetic diversity and ecosystem services (Morales et al., 2017; Vanbergen et al., 2018; Veron et al., 2018).

*Amegilla pulchra* (Smith, 1854) is a blue-banded bee common to the eastern cost of Australia. Its distribution range has recently increased dramatically over the past decade, after being introduced to several Pacific islands such as New Caledonia, Fiji and French Polynesia (Groom et al., 2017; Groutsch et al., 2019; Zakardjian et al., 2023b). Incidental introductions may have occurred with the maritime transportation of non-native ornamental plants and crops and seem to follow the main roads (Zakardjian et al., 2023b), suggesting that its spread may be facilitated through terrestrial transportation of alien plants as well. The species occasionally visits solanaceous plants using head-banging technique (sometimes referred to as buzz-pollination due to similarities), which makes it valuable for the pollination of crops such as tomatoes or eggplants (Groutsch et al., 2019; Hogendoorn et al., 2007; Udayakumar et al., 2023). Therefore, the possibility that *A. pulchra* may have been subject to deliberate introduction events cannot be ruled out. Most species introductions occur accidentally and *A. pulchra* is likely no exception to the rule. The species was first detected in urbanized environments of New Caledonia where it is now commonly found (Zakardjian et al., 2020), which makes it all the more likely to keep being spread.

We modelled the current invasion risks of a *Amegilla pulchra* accounting for the species climate niche and propagule pressure. To avoid the above-mentioned limitations, we used a modelling approach that performs well on small samples, accounting for non-equilibrium and assessing model spatial transferability. This approach enabled us to overcome the difficulties underlying IAS distribution modelling in order to (a) reliably identify the drivers of invasion and (b) identify the areas at risk at the global scale and provide robust recommendations for surveillance prioritization. Since we expect different effect of environmental change between the continent (corresponding to the native region) and islands (i.e. non-native regions), (c) we further compare the change in invasion risk between the native and the non-native regions.

## 2. Methods

*Overview* – we used an ecological niche modelling approach with a technique adapted to small samples (ensemble of small models), using a highly performing algorithm (RandomForest down-sampled), correcting for sample bias (coping with non-equilibrium assumptions), with a spatial partitioning for model evaluation (to assess spatial transferability), with a robust predictor selection process and accounting for anthropogenic factors and high-resolution global climate data that are appropriate for small islands.

*Occurrence data* – We used verified occurrence data from multiple sources from both the native (Australia) and non-native regions (Pacific islands), as recommended to best inform the models of the environmental conditions already filled by the species (Hui, 2022). We selected data points from GBIF (www.gbif.org), keeping data from the CSIRO Bee Trap Surveys, the research programme of Apoidea and iNaturalist research grade. We completed with a citizen science survey organised in New Caledonia by the French Institute of Research and Development (IRD). First broadcasted towards beekeepers, the programme has rapidly spread to the public and is now animated through a Facebook page and an email address. The program has been running for five years and has provided important data that are systematically validated through photo identification, *Amegilla pulchra* being easily distinguishable on picture from all other wild bees of New Caledonia. For the native area, we excluded observations with fuzzy coordinates or those based on pictures only (due to possible misidentification issues with closely related species such as *Amegilla chlorocyanea*). This results in an initial sample of unique point coordinates of 200 points. To avoid pseudo-replication, we resampled one point per pixel of environmental data at 1 km resolution (i.e. a process of spatial data thinning; Steen et al. 2021), resulting in a final sample of 102 points (42 from the native range and 60 from the non-native range).

*Climate data* – We used 19 bioclimatic variables (description available at https://www.worldclim.org/data/bioclim.html; see also Booth et al., 2014) at 30 arc seconds (approximately 1 km) resolution for the current and future (2070) climate from CHELSA version 1.2 (Karger et al., 2017). Predictions of climate suitability, as well as invasion risks can be highly sensitive to the choice of climate data source (Dubos et al., 2023a, 2022a). Besides CHELSA, Worldclim is the most commonly used global database available at 1 km resolution with multiple future projections available. However, Worldclim for less reliable when applied to small islands (IPCC, 2021), thus motivating us in using CHELSA only. The latter is based on statistical downscaling for temperature, and precipitation estimations incorporate orographic factors (i.e., wind fields, valley exposition, boundary layer height; Karger et al. 2017) and is the most appropriate for our study region.

For future projections, we used three Global Circulation Models (GCMs; i.e., BCC-CSM1-1, MIROC5, and HadGEM2-AO) and two greenhouse gas emission scenarios (one optimistic SSP2 and the most pessimistic SSP5) to consider a wide panel of possible invasion risk in 2070.

*Soil data* – *Amegilla pulchra* is a ground-nesting species. We hypothesized that soil density at the surface may be used as a proxy for the species’ capacity to dig its nest in the soil. We used surface bulk density data from the SRIC-WISE Harmonized global soil profile database (Batjes, 2009).

*Factors of spread* – To account for the risk of introduction and spread of the species, we used anthropogenic predictors which we assumed to be related to propagule pressure and spread. The species has been introduced accidentally through transportation of goods, notably horticultural exports. We used three types of drivers of introduction, propagule pressure and spread as proxies:

- distance to port and airports as a proxy for introduction risk. We obtained port data from the World Port Index (https://msi.nga.mil/Publications/WPI, accessed August 2024) and airport data from the OpenFlights Airport database (https://openflights.org/data.html, accessed December 2023).

- distance to main roads as an indicator of potential spread facilitation (Lanner et al., 2022), because the species is frequently observed on alien plants along roads. We computed the distance from roads using the Global Roads Inventory Project (GRIP4) dataset (Meijer et al., 2018). We selected the first two categories of road size (highways, primary roads).

- human population density as a proxy for both introduction and expansion risks, since the species is regularly observed foraging on ornamental plants (Zakardjian et al., 2020). We obtained human population density data for the current period and all four future scenarios from Gao (2020) at a resolution of 1 km².

For distance to ports, airports and distance to roads, we could not include future scenarios of change as such data are unavailable, so our future projections do not account for potential changes in these predictors.

*Background selection* – It is recommended to use a restricted background to mitigate the effect of sample bias, follow the hypotheses of past dispersal and account for the Equilibrium hypothesis (Acevedo et al., 2012; Dubos et al., 2022c; Vollering et al., 2019). We defined the background as a rectangle of minimal surface that includes all presence points, which we extend by 10 % in each cardinal direction. After model training on this background extent, we projected the predictions of invasion risk on a global map, excluding cold temperate climates from both hemispheres (latitude > 50°). For future projections, we used the restricted background for our measurements of environmental change effects (see *Quantifying the effect of environmental change* section), because the global maps may be more uncertain (as a result of spatial and temporal extrapolation).

*Pseudo-absence selection and sample bias correction* – We generated 15,000 pseudo-absences with the randomPoints function of the dismo package. This number represents a fair trade-off between model reliability and computation time (Valavi et al., 2021). We generated three sets of pseudo-absence for each block-cross validation run (see *Model evaluation* section for more details about spatial blocks). We applied a sample bias correction technique consisting in a selection of pseudo-absence points that imitates the spatial bias of presence points (Phillips et al., 2009b). To do so, we used a null geographic model, which is the most appropriate approach for species that are still spreading and have not reached equilibrium, because it downplays the areas that have not been invaded yet and therefore mitigates the effect of non-equilibrium. We used the null geographic model as a probability weight for pseudo-absence selection.

*Predictor selection* – The selection of environmental predictors is an unresolved issue in SDM (Leroy, 2023). To best fit with the latest methodological recommendations (Araújo et al., 2019; Gábor et al., 2019; Sillero et al., 2021; Sillero and Barbosa, 2021), we performed a predictor selection process in five steps: (1) removing irrelevant predictors (Araújo et al., 2019), (2) removing collinear predictors (Dormann et al., 2013), (3) keeping predictors with the clearest causal relationship with species presence and spread (Dubos et al., 2022d; Fourcade et al., 2018; Hui, 2022), (4) statistical selection based on relative importance (Bellard et al., 2016b; Thuiller et al., 2009), (5) consideration of potential interactive effects (Gábor et al., 2019). (1) We discarded bio3 (isothermality) because the relationship with species presence was unclear. We kept the remaining 18 because they all may explain species distribution (mostly through direct effects on temperature and water balance, or indirect effects on ecosystem productivity and food availability; (Dubos et al., 2018b). (2) For each climate data source and for each species-specific background (details below), we selected one predictor variable per group of inter-correlated variables to avoid collinearity (Pearson’s r > 0.7; (Dormann et al., 2013) using the removeCollinearity function of the virtualspecies R package (Leroy et al., 2016). (3) When mean values were collinear with extremes, we selected the variables representing extreme conditions (e.g., warmest/driest condition of a given period) because these are more likely to drive mortality and local extirpation and be causally related to the species’ presence (Maxwell et al., 2019; Mazzotti et al., 2016). (4) We excluded the predictors with the lowest relative importance (assessed by the GINI index of the RandomForest R package). Relative importance was assessed with centred and scaled predictors. (5) To consider potential interactive effects while accounting for propagule pressure, we kept a final set of two predictors related to temperature, two related to precipitation, and two related to anthropogenic introduction and spread.

*Modelling technique* – We modelled and projected climatic niches using a single highly performing tuned algorithm (Valavi et al., 2023, 2021), Random Forest down-sampled (hereafter RF down-sampled, i.e. RF parametrised to deal with a large number of background samples and few presence records; Prasad et al. 2006). We set RF down-sampled to run for 1000 bootstrap samples/trees. We used an Ensemble of Small Models approach (ESM, Breiner et al., 2015), which performs well with single-algorithms (Breiner et al., 2018). This approach is adapted to small samples, allows to account for multiple environmental factors (therefore maximizing the explanatory power), while being parsimonious (thus limiting the possibility of overfitting). It consists in running combinations of bivariate models. In the case of an invasive species, it is important to account for factors of spread. We adapted the ESM approach to predicting invasion risks by systematically including two factors of spread with the combinations of two climate predictors. Since our final set of predictors included four climate variables and two variables of anthropogenic introduction and spread (see *Predictor selection* section), this results in six model replicates (i.e. 6 combinations of two climate predictors + two factors of spread). After discarding poorly performing models (see *Model evaluation* section), we averaged all projections across all replicates.

*Model evaluation* – We used block-cross validation approach which better assesses model transferability compared to random partitioning (Valavi et al., 2023, 2019). We spatially partitioned our data into four square blocks of 1000 km (Fig. S1). We assessed model spatial transferability with the Continuous Boyce Index (CBI; Hirzel et al., 2006), because other discrimination metrics such as AUC and TSS require absence data and may be misleading with biased presence-only data (Dubos et al., 2022d; Leroy et al., 2018). An index of 1 suggests that suitability models predicted perfectly the presence points, a zero value suggests that models are no better than random and a negative value implies a counter prediction. We discarded all models with a Boyce index < 0.1. We present projections of the mean suitability obtained across all fairly transferable models (total number of models: 4 block-cross validation runs × 3 pseudo-absence runs × 6 ensemble of small models = 72; for future projections: 72 × 2 scenarios × 3 GCMs = 432 projections).

*Quantifying the effect of environmental change* – We quantified the projected change in environmental suitability (1) at the presence points, at the presence points of (1a) the native and (1b) invaded area separately, (2) across the model training background. At the presence points, we computed the difference between predictions of future and current invasion risk to represent the effect of environmental change on current populations. When considering the background, we computed the difference between the averaged scores of invasion risk for current and future predictions, expressed in percentage of change across the background. This metric is similar to the Species Range Change (SRC, Buisson et al., 2010; Dubos et al., 2022a) and provide information on the overall direction and magnitude of the change across the region.

*Quantifying climate niche overlap –* We assessed the level of similarity between the climate conditions of the native and the non-native regions. We computed a Principal Component Analysis between the selected climate predictors of the native and non-native ranges (bio2, bio10, bio18 and bio19) following Broennimann et al. (2012). We quantified niche overlap with Schoener’s D overlap index and visualized both climate niches (as represented by two principal components) using the *ecospat.plot.niche.dyn* function of the ecospat R package (Di Cola et al., 2017).

*Identifying priority for surveillance –* We ranked global ports by invasion risk and maritime traffic to provide guidelines for proactive surveillance. We accounted for environmental suitability and factors of spread using model projections of invasion risks of obtained from the previous steps. For each port, we averaged predictions of invasion risk within the surrounding 100 km of the port. We chose a buffer of 100 km radius because we found a high invasion risk until this distance approximately (see Results section). Beyond this distance, predictions of invasion risk were intermediate, but most likely due to ports located in the neighbouring islands. Since the species spread is facilitated by transportation of goods, we also accounted for maritime traffic using global data from the Global Maritime Traffic maintained by MapLarge (https://globalmaritimetraffic.org/). We produced an index of surveillance priority for every port documented in the World Port Index between -40 and 40° Latitude for all countries that are located around the Pacific Ocean and the Indo-Pacific region. The index is the product of invasion risk and number of vessels per year, both centred and scaled.

## 3. Results

The transferability of our model set was highly variable. We discarded 33 poorly performing models out of 72 (mean CBI = 0.16 ± 0.50 SD). Based on relative importance, assumptions of causal relationships and consideration of interactive effects, we selected four climate predictors (Bio2, Bio10, Bio18, Bio19) and two predictors of propagule pressure (human population density and distance to closest port; Fig. S2). Although the explanatory power of Bio10 was lower, we decided to include this predictor into the final set to account for potential interactive effects between precipitation and temperature. The response to soil density was meaningful (the species tends to avoid the densest soils), but the explanatory power of the predictor was poor nonetheless (Fig. S3) and we chose not to include it in final models. The most important predictor was Bio2 (temperature diurnal range), followed by the two factors of propagule pressure (human population density and distance to ports). *Amegilla pulchra* avoids regions with high daily temperature variation (Bio2), also avoids the warmest and driest regions (mean summer temperature > 29 °C, summer precipitation < 2000 mm, winter precipitation < 500 mm). The species is absent from unpopulated regions, and the invasion risk decreases as the distance from a port increases (very high until 100 km, null beyond 400 km; Fig. S4).

The predicted current invasion risk is particularly high in the Indo-Pacific islands, Southeast Asia, eastern China, South Korea, Japan, Florida and the Caribbean (Fig. 1).

**Fig. 1.**
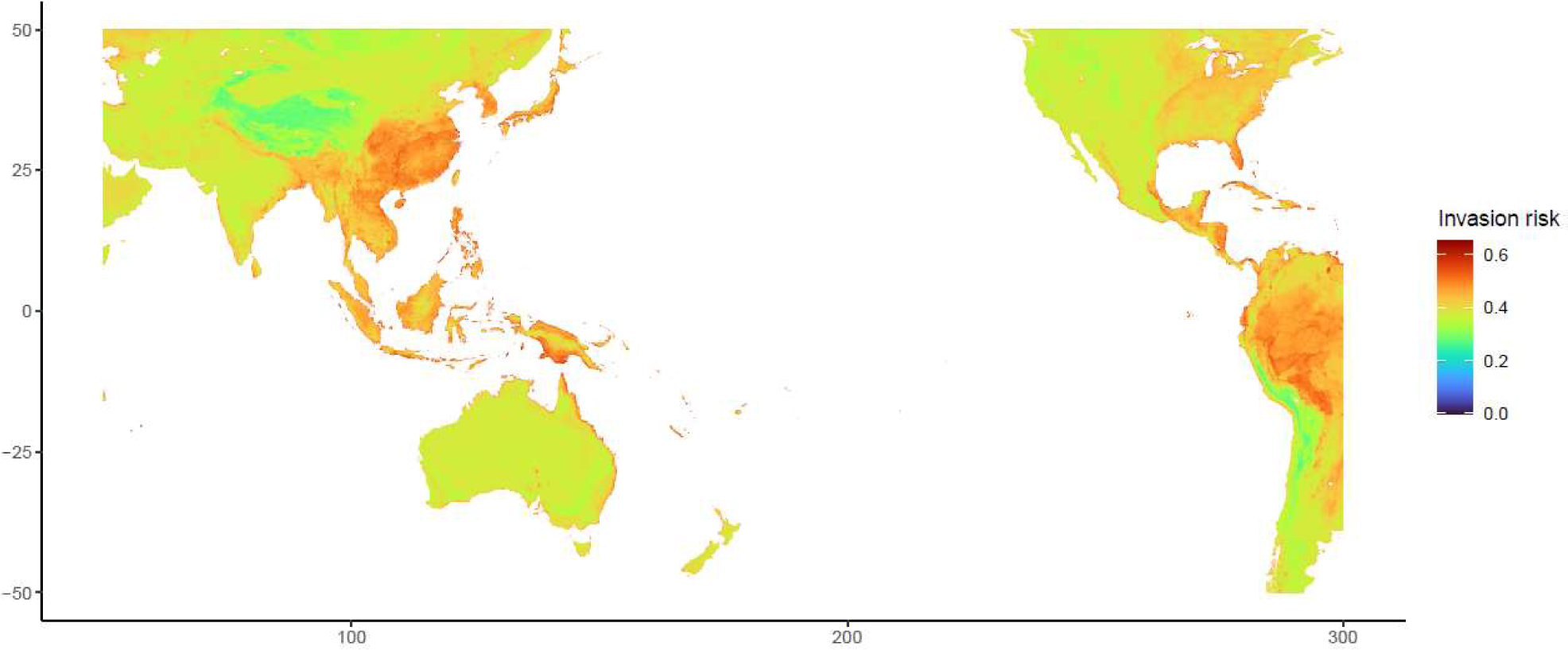
Current global invasion risk of *Amegilla pulchra*. Predicted values were obtained from Species Distribution Models and included climate and anthropogenic factors. A high-resolution version is available in the online material.

Invasion risks will vary in either direction in the future, depending on the region (Fig. 2, 3). At the level of the presence points, invasion risk will increase by 20.7 % ± 29 SD (Fig. 2). The predicted change is significantly more variable in the native range (+19.9 % ± 39.2 SD) than in the non-native one (+21.3 % ± 21.3 SD; Fig. 3; variance test for the six future projections: F-statistics ranging between 0.17 and 0.25, p-values ranging between 10^-6^ and 10^-9^). Although the increase in invasion risk was higher in average in the non-native range, the difference with the native range was not significant (ranges of Wilcoxon tests for the six projections: W = [947-1083], p-values = [0.54-0.95]). Across the whole background (eastern Australia and the southern Pacific islands), the invasion risk will decrease by 78.2 % ± 1.2 SD on average across scenarios and GCMs.

**Fig. 2.**
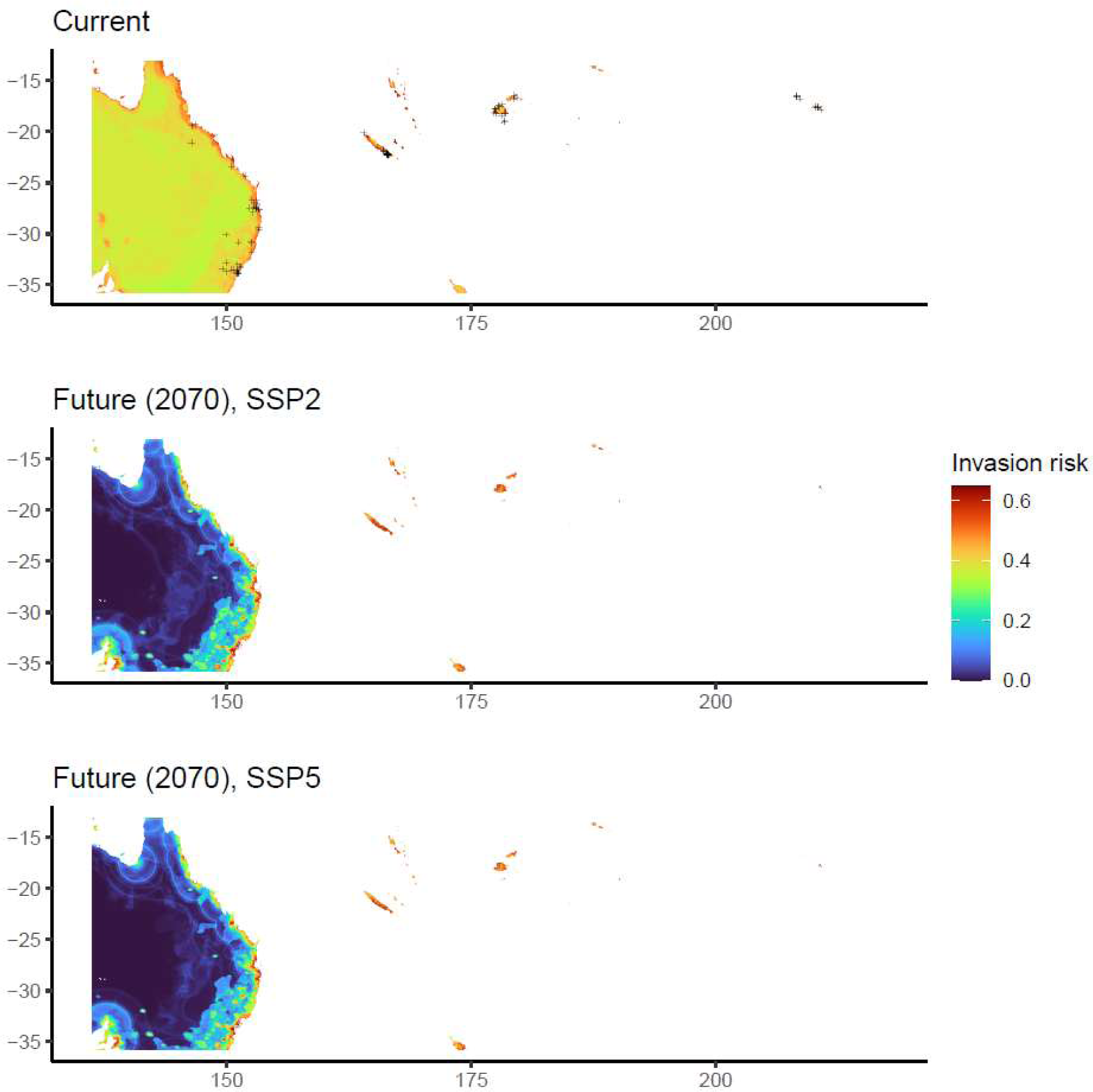
Projected current and future (2070) invasion risk of *Amegilla pulchra* in eastern Australia and Pacific islands according to two scenarios of the IPCC (top: current; middle: SSP2; bottom: SSP5). Presence points are represented with ‘+’.

**Fig. 3.**
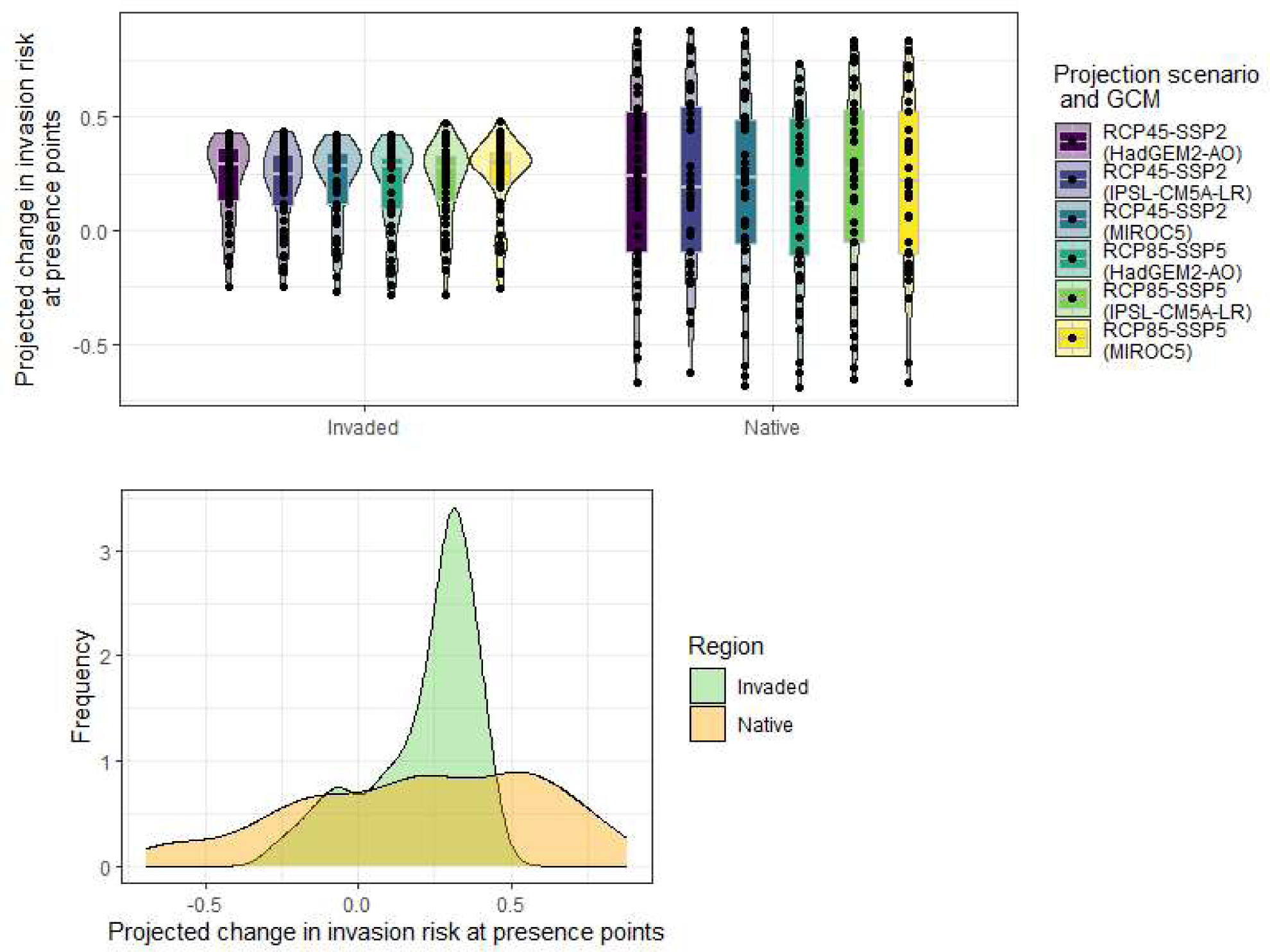
Predicted change in invasion risk by 2070 for *Amegilla pulchra* at current locations of the non-native (invaded) and native range across two scenarios and three Global Circulation Models (GCMs). Top panel: Scenario-GCM specific violin plots; Bottom panel: frequency distribution. The predicted change is the difference between future and current suitability expressed as a ratio (ranging between -1 and 1). Positive values mean a predicted increase in invasion risk, zero means no change. Boxes are composed of the first decile, the first quartile, the median, the third quartile and the ninth decile.

The introduction of *Amegilla pulchra* has largely led to the extension of its climate niche (Fig. 5, S6; Schoener’s D = 0.11). Some climate conditions of the species native range (green + light green + blue) are available in the non-native area (red contour) but still unoccupied (green), suggesting a potential for further spread.

**Fig. 4.**
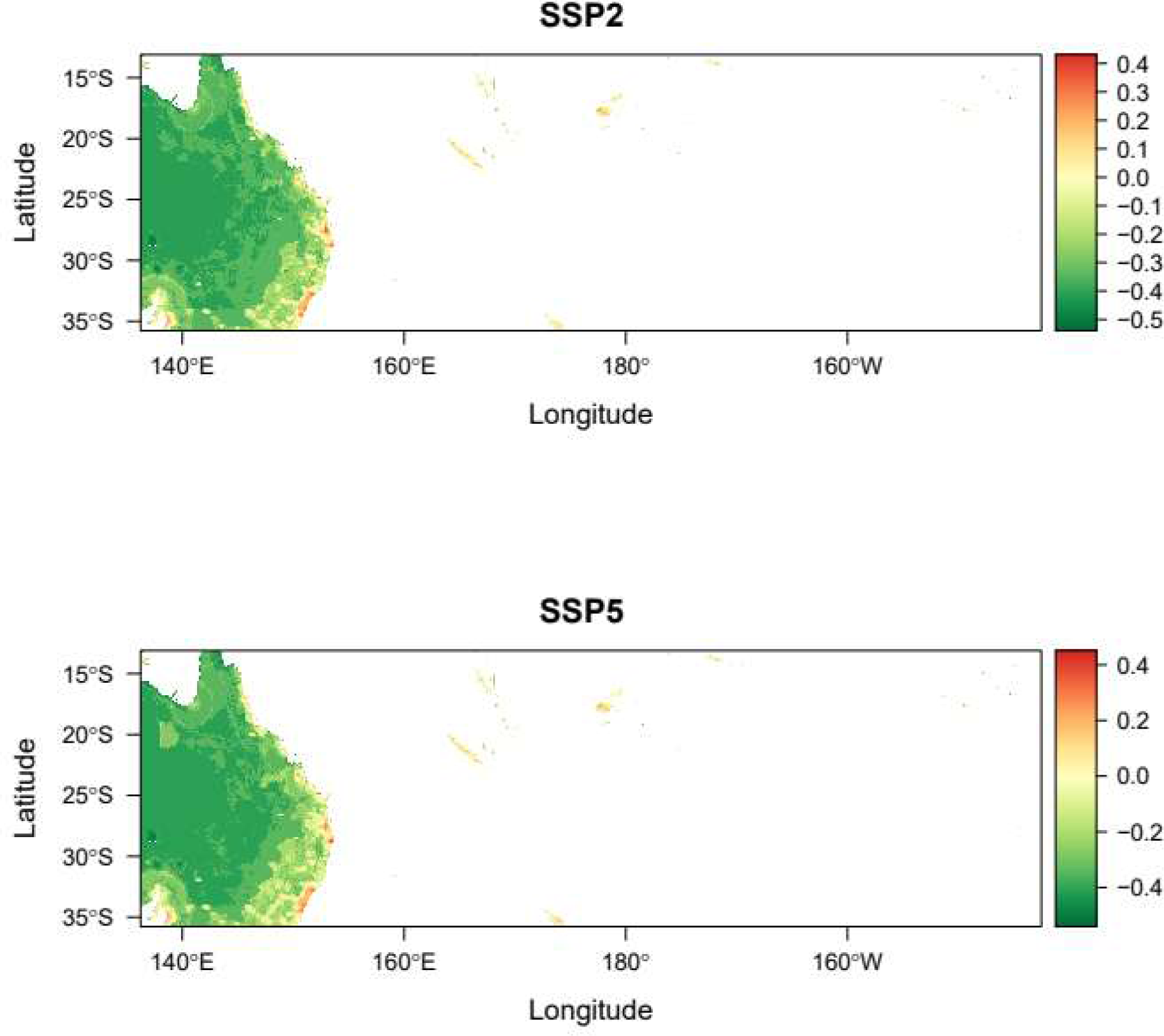
Projected temporal change in invasion risk for *Amegilla pulchra* by 2070 according to two scenarios (red: increasing risk; blue: decreasing risk).

**Fig. 5.**
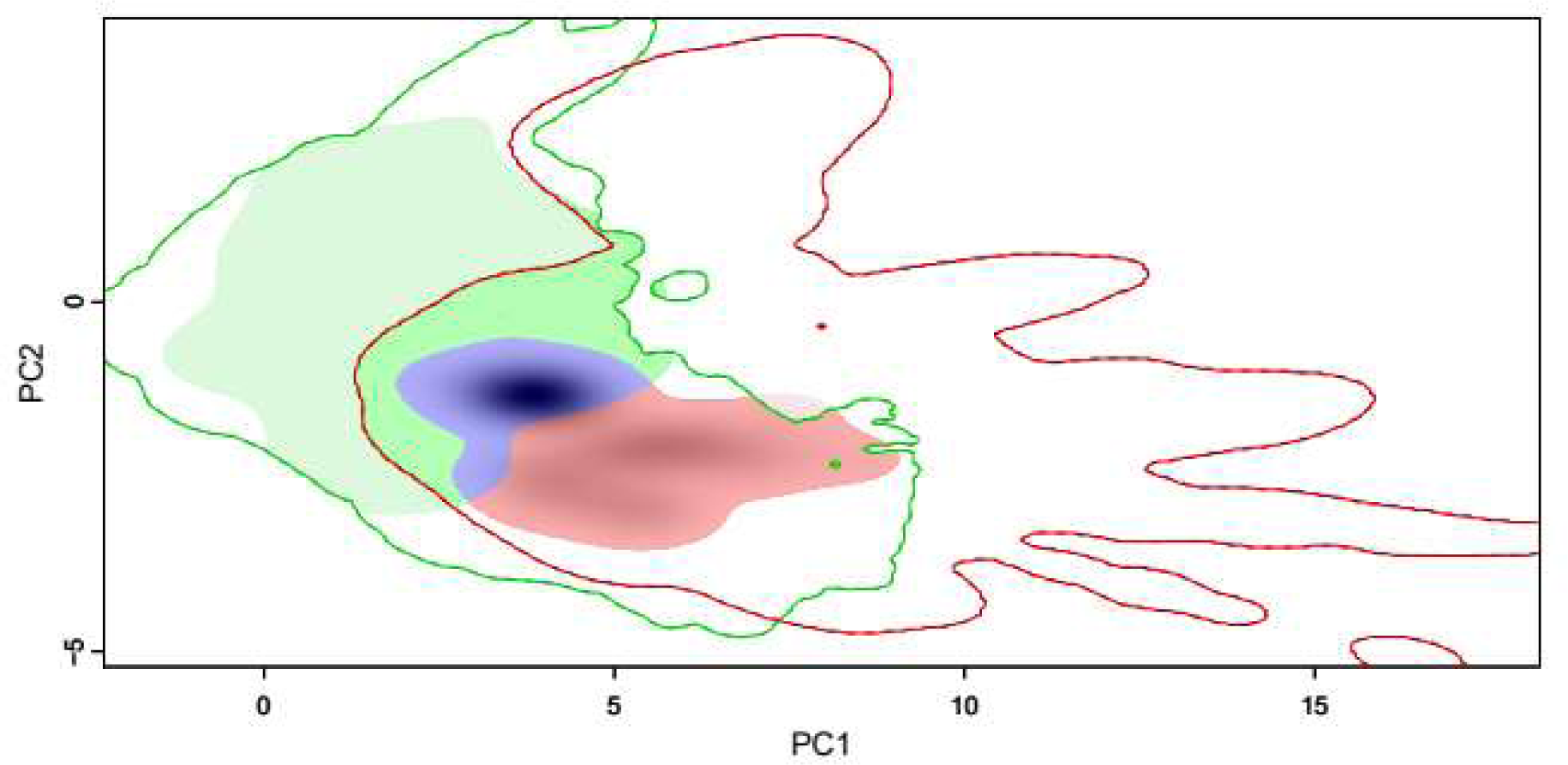
Realized climate niche of *Amegilla pulchra* obtained from Principal Component Analysis (four climate predictors: bio2, bio10, bio18 and bio19; see figure S6). Green: climate niche of the native range; red: niche expansion (conditions of the non-native range); blue: niche stability (i.e. overlapping conditions). The solid contour lines indicate the extent of environmental conditions that are available respectively in the native and invaded ranges. The shades of black represent the density of occurrence points at a given set of principal component (PC) score.

The ports with the highest priority for surveillance under current conditions were mainly located in Singapore and China (1^st^: Keppel East-Singapore, 2^nd^: Jurong Island (Singapore), 3^rd^: Zhoushan (China); Table S1).

## 4. Discussion

With a robust modelling approach, we predicted the invasion risk of a rapidly expanding solitary bee introduced outside of its native range a decade ago. We found a high potential of spread world-wide. The risk will vary in the future, decreasing in most of the native region but increasing in human-dense areas and throughout the non-native regions. Invasion risks will consistently increase in islands of the non-native range, highlighting a risk for insular ecosystems.

### 4.1 Drivers of Amegilla pulchra distribution

Temperature – *A. pulchra* is mostly found in areas with low diurnal variation in temperature, and avoids the hottest regions. This may be related to its foraging behaviour (Elias et al., 2017). Similar to other crop pollinators, the species seems to be active within a given temperature range (Jaboor et al., 2022). Those thermal conditions may also characterize the suitable temperatures for larval development (Lanner et al., 2022). The set of predictors we used (at the scale of the month or quarter) do not allow us to determine a minimum nor maximum temperature threshold for activity or larval development. Determining such threshold should be further investigated with experimental design at finer temporal resolutions. Ectotherms are generally little capable of handling large variation in thermal conditions at short temporal scales, which might also be the case for *A. pulchra.* The species seems to prefer sub-tropical to tropical environments and coastal environments in its native area. For now, most of the invaded areas are located on islands near the coasts, where the climate is similar to its native region. Our projections showed that continental environment are suitable too, especially in the Korean peninsula, China and Japan. We strongly recommend to closely monitor the spread of the species and assess its ability to adapt effectively to this type of environment.

Precipitation – The species is currently covering a large range of precipitation regimes, only avoiding very dry environments. This is consistent with the current distribution of the species, which is absent from the dry regions of central Australia. This could be related to physical aspects and the ground-nesting habits of the species: dry and compacted soils may hamper *Amegilla* bees for digging. This is also consistent with the findings of a high consistency in evaporative water loss across gradients of temperatures (Tomlinson et al., 2015), which may explain why the species is distributed from south to north of the Australian east coast. In addition to direct physiological constrains, the avoidance of dry regions may be related to the availability of host plants, which presumably require a minimum amount of yearly precipitation. The diet of the species is poorly documented. As far as we know, *A. pulchra* tends to prefer alien plants in invaded regions (Zakardjian et al. 2020). But Apidae *spp*. are also very generalist (long-tongues species) and may be able to collect nectar and pollen on a variety of flowering species. A better knowledge of the preferential host plant of the species could certainly meliorate our understanding of the species ecology and capacity to establish in new environments.

Soil characteristics – Although *Amegilla pulchra* is a ground-nesting species, soil density was a poor predictor of its current distribution. One possible explanation is that the density and nature of the soil are not discriminant for the nesting ecology of the species. Alternatively, this may be due to a high plasticity in nesting strategies. Indeed, while most of ground-nesting bees are strictly restricted to underground cavities for their brood, the species was observed nesting in rock cavities or trunks in New Caledonia (pers. obs. M.Z.) suggesting an important plasticity, therefore a high potential for further invasion. We remain very cautious with this information and this kind plasticity should be thoroughly verified in the future as it is not so common among bees. A notable observation which resulted from the citizen science program is that, in invaded regions such as New Caledonia, the species has never been observed in ultramafic soils (Zakardjian et al., 2023b), a parameter which might be accounted for in future studies (provided the data is available throughout the whole study region). To date, the species is restricted to calcareous soils in New Caledonia. Ultramafic substrates are characterized by a nutrient deficiency and high concentrations of heavy metals and metalloids. The first parameter constrains plant growth, leading to high levels of endemism in ultramafic environments and the presence of hyper-accumulating species (Lannuzel et al., 2021). Such a floristic composition could be non-attractive perhaps repellent to exotic flower-visiting insects. *Amegilla pulchra* may avoid ultramafic soils either as a result of lack of host plants, or due to high concentrations of nickel acting as a chemical barrier, or as a physical barrier preventing from building nests (Zakardjian et al., 2023b). We encourage the pursue of our citizen science programme towards the blue banded bee in invaded regions and shed a light on the nesting behaviour on the species.

Human density and distance from ports – Similar to most introduced species, *Amegilla pulchra* is found is human-populated regions. Individuals may have been incidentally spread through maritime traffic of ornamental plants, as well as crops, which would explain the higher presence within approximately 40 km around ports. Notwithstanding the possibility that this result may have been related to an observer bias, we cannot exclude that the species may also have been deliberately introduced for its efficiency in the pollination some solenaceae, which would contribute to its proximity to human infrastructures.

Proximity to large roads – Although the species’ spread is likely facilitated by road traffic (Zakardjian et al., 2023b), the distance from roads did not explain well the current distribution of the species. This is presumably due to the absence of road categorized into the largest two categories of road size in small islands (Meijer et al., 2018). This suggests that the spread might be facilitated not only by commercial traffic, but also by private individual transportation of goods.

Comparison native *versus* non-native climate niche – We found an important niche extension in the non-native area. The climate niche of the non-native range (red area on figure 5) can be found in the native region where the species is not present (green contour, Fig. 5). The absence of the species in these environments of the native region may be explained by biotic interactions (e.g. competition with other *Amegilla* species; Leijs et al., 2017). This suggests that the species has extended its realized niche as a result of ecological release (Fieldsend et al., 2021). Reversely, the climate niche of the native range (green) is represented in the non-native region (red contour) but the species is not present yet, which suggests that there is room for further spread towards environments that are known to be suitable to the species.

### 4.2 Predicted effect of climate change

Future climate change might induce range expansions in some Australian bee species due to the availability of cooler conditions in the altitudes (Dew et al., 2019). Our predictions suggest that this will not be the case for *Amegilla pulchra* in the native range. We found a highly contrasting change in environmental suitability throughout eastern Australia. Given the wide latitudinal gradient characterizing the species distribution, one might explain this variability by a differential sensitivity to climate warming through latitudes at the population scale (Deutsch et al., 2008; Dubos et al., 2018a; Kazenel et al., 2024). However, our predictions indicate a negative change in invasion risk either in the north or the south of the native range. The regions of increasing risk are located in highly human-dense areas, suggesting that climate change might be detrimental to the species but its spread will keep being facilitated by human activity. This result is consistent with another Australian bee species (*Ceratina australensis*), which is predicted to increase in urban environments (Dew et al., 2019). However, such prediction is more likely driven by the predicted increase in human population rather than climate suitability.

Temporal changes in both temperature and precipitation may induce physiological stress, either at the larval or adult stage (Kazenel et al., 2024). Higher temperatures induce an increase in evaporative water loss in *Amegilla* spp. (Tomlinson et al., 2015). Detrimental effects might also derive from indirect effects on plants and their interaction with their pollinator (Scaven and Rafferty, 2013). Climate change and subsequent distribution shifts could drive potentially important spatial mismatches between crops and their pollinators, as it is the case for apples (Marshall et al., 2023). The mismatch could also be temporal, with differential phenological shifts between the pollinator and its host plants (Weaver and Mallinger, 2022).

Despite the overall decrease in invasion risk in the native range, we predict a more consistent positive change in invasion risk in the non-native range. Such difference may be explained by a stronger contrasts in human density in the native range. The difference may also be explained by the strong mismatch between the realized niche of the native and the non-native range, as shown by our climate niche analysis (Fig. 5). So far, the species established in Pacific Islands, characterized by higher precipitation regimes and more consistent temperatures. The success of invasive bees may be explained by larger thermal tolerance and higher dessication resistance, as it is the case in Fiji (Da Silva et al., 2021). In the future, the potential spread of an invasive bee such as *Amegilla pulchra* may be detrimental to local pollinator species as result of better resilience to a changing environment (Da Silva et al., 2021). The effects on local systems may be aggravated through the facilitation of spread of alien plants or the introduction of new parasites (Meeus et al., 2011; Zakardjian et al., 2023a).

### 4.3 Limitations

Our modelling approach could characterize the climate niche of the species with the available high-resolution data, and take into account factors of introduction and spread. We applied the latest recommendations regarding the modelling of a species that is still spreading and assessment of spatial transferability of our predictions. One limitation of our approach might be the misidentification of *Amegilla pulchra* in the native range (confusion is unlikely in the non-native range), which is often confounded with other *Amegilla* species despite photographic validation on online platforms. However, such limitation may not have led to erroneous assessments, because the other *Amegilla* species share a similar ecological niche. A potential avenue for further studies is to assess the effect of climate on species activity and reproduction and test for differences along its distribution range, and between the native and invaded areas. Another limitation is related to the small sample size and incompleteness of our input data. Projections are likely to change with updates of the known distribution of the species. However, we used the available tools and data to mitigate this effect and our models still could identify high invasion risks in non-invaded regions.

### 4.4 Management recommendations

The eradication of an invasive species is challenging and costly, especially on the continent (Diagne et al., 2021; Pearson, 2024). In regions where the species has not been introduced yet, preventing species from arriving is by far the most effective approach (Cuthbert et al., 2022; Pearson, 2024). We identified priority areas for surveillance efforts for prevention in areas that are not invaded yet. In ports identified as highly at risk of invasion and the surrounding regions, local stakeholders should be informed and trained to species identification.

In their native range, *Amegilla* spp. are recognized as efficient pollinators of solanaceae which require buzz or head-banging pollination. Therefore, several species have been used in greenhouses to enhance crop yield such as tomatoes and eggplants (Hogendoorn et al., 2006; Udayakumar et al., 2021). This element raises concerns as it reminds the case of *Bombus* spp., that perform buzz pollination, and were voluntarily introduced worldwide for promoting greenhouse tomato pollination (Aizen et al., 2019). Now, *Bombus* spp. – such as other managed alien bee species – are recognized as a major driver of native pollinator declines and plant-pollinator interaction disruptions (Morales et al., 2013; Aizen et al., 2019; Geslin et al., 2023). Out of principle of precaution, the use of the species should be prohibited.

In regions where the species has already been spread, our future projections help identifying priority areas for control and mitigation measures. *Amegilla pulchra* is commonly found on alien plants (Zakardjian et al., 2023b). In the non-native regions, habitat restoration may be a promising avenue for control and spread mitigation. This would benefit local ecosystems through the extirpation of alien plants and the promotion of local communities.

Climate change will alter invasion risks in the future. For instance, in New Caledonia the risk will decrease in the north but increase in the south (Fig. 4); On the island of Tahiti, the risk will increase in Nui (North of the island) and decrease in Iti (South). Priority areas for management will need to shift accordingly in the future for cost-effectiveness.

### 4.5 Conclusions

*Amegilla pulchra* may keep spreading in already invaded areas and colonize further regions. Since the species may be introduced deliberately, public awareness might be the best strategy to prevent further introductions and mitigate the potential effect of the species on local systems. We call the need to closely monitor the spread of *Amegilla pulchra*, promote habitat restoration in already invaded areas and diversify crop pollinators in order to preserve island ecosystems.

## Acknowledgements

We sincerely thank all participants of our citizen science programs for the collection of occurrence data. We gratefully acknowledge Prisca Mahé, Thomas Cochenille, and Valentin Mitran for their invaluable contribution in collecting occurrence data for Amegilla pulchra during research programs in New Caledonia. We also extend our thanks to the New Caledonian Centre de Promotion de l’Apiculture and Province Sud for their support in the dissemination of the citizen science program.

## CRediT authorship contribution statement

**Nicolas Dubos**: Conceptualization, Data Curation, Formal analysis, Methodology, Writing – original draft. **Benoit Geslin**: Conceptualization, Investigation, Validation, Writing – review and editing. **Hervé Jourdan**: Investigation, Writing – review and editing. **David Renault**: Writing – review and editing. **Marie Zakardjian**: Conceptualization, Data Curation, Writing – review and editing.

## Declaration of competing interest

The authors declare that they have no known competing financial interests or personal relationships that could have appeared to influence the work reported in this paper.

## Notes

### Competing Interest Statement

The authors have declared no competing interest.

